# Electrophysiological Characteristics of Epidural Spinal Signals in Preclinical Models of Spinal Cord Stimulation

**DOI:** 10.1101/2025.09.03.674002

**Authors:** Kimberley Ladner, Eline M. Versantvoort, Marjolein E. G. Thijssen, Dave Mugan, Stuart N. Baker, Alexander Kraskov, Quoc C. Vuong, Mahima Sharma, Marom Bikson, Birte E. Dietz, Stefano Palmisani, Ilona Obara

**Affiliations:** School of Pharmacy and Translational and Clinical Research Institute, Newcastle University, Newcastle-upon-Tyne, UK; Saluda Medical Europe Ltd, Harrogate, UK; Biosciences Institute, Newcastle University, Newcastle-upon-Tyne, UK; School of Psychology, Newcastle University, Newcastle-upon-Tyne, UK; Neural Engineering Laboratory, Department of Biomedical Engineering, The City College of the City University of New York, City College Center for Discovery and Innovation, USA; Guy’s and St. Thomas’ NHS Foundation Trust, London, UK

**Keywords:** evoked compound action potentials, doublets, evoked synaptic activity potentials, spinal cord stimulation, rat, macaque

## Abstract

**Objectives:** Epidural stimulation of the spinal cord evokes distinct electrophysiological responses that can be recorded epidurally. Here, we characterized evoked compound action potentials (ECAPs), doublets (secondary or tertiary ECAPs, likely of different physiological origin than primary ECAPs), evoked synaptic activity potentials (ESAPs), and electromyographic (EMG) signals in preclinical models. Our objective was to clarify the features and distinct physiological origins of these signals, in order to advance mechanistic studies and support clinical applications of spinal cord stimulation (SCS) therapy.

**Materials and Methods:** Adult male Sprague-Dawley rats (300-440 g) were implanted with two epidural leads (caudal and rostral; each with eight electrodes) and received monopolar, biphasic stimulation (200 μs pulse width) at 2 and 50 Hz, with current increased stepwise to motor threshold. Rhesus macaques (11.5 and 10.2 kg) were implanted with a single 12-electrode epidural lead and stimulated using either tripolar, triphasic pulses at 10 Hz (100 μs) or tripolar, biphasic pulses at 3 Hz (80 μs) up to 3×ECAP threshold. Recordings were taken from non-stimulating electrodes.

**Results:** ECAPs and EMG signals were recorded across multiple spinal segments in both rats (L1-T7) and macaques (L2-T11). Doublets presented as complex waveforms with multiple negative peaks, two in rats and three in macaques, likely representing distinct ECAPs at T11-T6 in a rat and L1-T11 in macaques. ESAPs, detectable in rats, showed anatomical specificity, over the L1/T13 vertebrae with peak responses at L1. Signal analysis included activation thresholds, amplitudes, latencies, and conduction velocities.

**Conclusions:** This study outlines electrophysiological signals evoked by SCS in terms of their waveform, recruitment thresholds, and putative physiological origins. We propose that, to the extent these signals reflect different aspects of spinal processing and may serve as biomarkers of dysregulated nociceptive pathways, as well as indicators of SCS efficacy or potential side effects.

## INTRODUCTION

Over the past decade, the technology to record electrophysiological signals from the epidural space in both humans and animal models during dorsal column (DC) stimulation has advanced spinal cord stimulation (SCS) therapy and research [1–5]. In particular, the recording of evoked compound action potentials (ECAPs), which reflect the synchronous activation of DC fibers, have enabled the development of a closed-loop SCS system [6–8]. ECAPs, when integrated into the closed-loop system, provide a real-time and pragmatic biomarker of spinal cord activation. This enables continuous monitoring of neural activation and dynamic adjustment of stimulation parameters to maintain DC activation, which has been linked to improved outcomes in pain patients [9–12]. Beyond their clinical utility, ECAP recordings also offer insights into the neurophysiological mechanisms underlying SCS, providing a direct and dynamic view of neural modulation during stimulation [13, 14].

Recent studies have identified additional electrophysiological signals recorded via epidural SCS electrodes that reflect neurophysiological responses to stimulation distinct from ECAPs [15, 16]. Sharma et al. (2023) identified a novel electrophysiological signal during epidural SCS in rats, termed evoked synaptic activity potentials (ESAPs), which reflect synaptic activity within dorsal horn neurons following DC activation [15]. ESAPs consist of several slow waveforms, most notably the S1 component, which appears after the triphasic ECAP signal (P1-N1-P2). The synaptic origin of ESAPs was validated by administering a glutamatergic synapse blocker (6-Cyano-7-nitroquinoxaline-2,3-dione; a selective competitive AMPA receptor antagonist) which significantly reduced the S1 wave without affecting the ECAP [15]. A separate fast evoked potential, referred here to as a secondary ECAP, or doublet, occurring ∼1-2 ms after the primary ECAP has been reported by Gmel et al. (2023) in response to SCS in humans. They proposed the involvement of post-synaptic dorsal column (PSDC) fibers, which may be activated either synaptically by primary afferents or directly by stimulation pulses [16]. These secondary responses exhibited distinct conduction velocities, suggesting they arise from a separate neural population than those contributing to the primary ECAP [16].

Herein, we discuss the possibility that these likely second-order signals propagate through one of several spinal tracts [17]. Finally, electromyographic (EMG) signals, recorded from epidurally placed electrodes across multiple species, represent muscular responses to epidural stimulation, arising either from a Hoffmann reflex or direct activation of motor neurons [18].

The novel evoked spinal potentials, like ESAPs and doublets, are compelling because they reflect neural processing distinct from ECAPs, offering complementary biomarkers for therapeutic engagement and mechanistic insight. However, epidural recordings during SCS are also susceptible to myogenic signals and various stimulation artifacts, and therefore rigorous signal processing is essential to accurately distinguish neurophysiological responses from myogenic or electrical noise [19, 20].

To advance the understanding of electrophysiological signals evoked by SCS, we analyzed experimental data from both rats and macaques and reviewed the existing literature on these signals. Our objectives were to systematically characterize signal features across species, discuss their potential as biomarkers and mechanistic indicators of SCS efficacy, and identify critical factors that influence their interpretation and applicability.

## MATERIALS AND METHODS

### Animals

#### Rat

Naïve adult male Sprague-Dawley rats (n = 18; 8-10 weeks; 300-440 g; Charles River Laboratories, Kent, UK) were acclimatized to the colony room for at least seven days after arrival and prior to the start of any experimental procedures. Rats were housed in groups of 3-4 animals per polyethene cage (Comparative Biology Centre, Newcastle University, Newcastle upon Tyne, UK). They were maintained on a 12-h day/night cycle (lights on at 6:00 am; lights off at 6:00 pm), and under controlled temperature (21°C) and humidity (55%). Standard rodent food and water was accessible *ad libitum*.

#### Macaque

Adult male rhesus macaques (n = 2; ∼7 years; 11.5 and 10.2 kg; Centre for Macaques, Salisbury, UK) were pair-housed in spacious pens exceeding UK and EU welfare standards, with access to natural daylight (Comparative Biology Centre, Newcastle University, Newcastle upon Tyne, UK). These animals were part of a separate, unrelated study involving unilateral corticospinal tract lesions, induced either by intracortical injections of endothelin-1 or by a thermocoagulation of the internal capsule. Water was available *ad libitum,* and food access was regulated to maintain motivation during the behavioral components of the primary study.

All experiments were conducted in accordance with the UK Animals (Scientific Procedures) Act 1986 under Home Office Project Licences P6694C943 and PP8237166, with local ethical approval from the Animal Welfare and Ethical Review Body (AWERB). This study adheres to the Animal Research: Reporting of In Vivo Experiments (ARRIVE) guidelines. Animal welfare was carefully monitored throughout the study, and all efforts were made to minimize suffering and reduce the number of animals used.

### Lead Implantation and SCS Delivery

#### Rat

General anesthesia was induced via a nose cone using 5% isoflurane in oxygen (2 L/min flow rate). After induction, urethane (1.5 g/kg, i.p.) was administered alongside glycopyrrolate (0.015 mg/kg, s.c.) and anesthesia was maintained with 0-0.5% isoflurane in oxygen (2 L/min flow rate). All rats were implanted with two custom-made, eight-electrode epidural SCS paddle leads (flexible polyamide printed-circuit design with gold-plated electrodes; overall dimensions: 1.25 mm width × 0.1 mm thickness; individual electrode size: 0.3 mm width × 1.0 mm length × 0.1 mm thickness). The leads were designed by Saluda Medical (Harrogate, UK) and fabricated by PCBWay (Hangzhou, China), and were placed in the dorsal epidural space. Each lead featured electrodes spaced 4 mm apart (center-to-center). Electrode placement was verified by X-ray imaging (Orange 1040HF, EcoRay, Seoul, South Korea) (Figure 1a). Implantation procedures followed protocols established in our previous studies [1, 10]. One lead was positioned caudally, targeting spinal segments L2-T12, while the second lead was placed rostrally, spanning T12/T11 to T6 (Figure 2a). Both leads were connected to a custom-designed Multi-Channel System MKII (Saluda Medical, Sydney, Australia) as described in our earlier work [1, 2, 10]. Monopolar, biphasic stimulation was delivered in a stepwise manner with increasing current up to visible motor threshold (observed as a contraction in back or leg muscles), using a pulse width of 200 μs, at frequencies of 2 (n = 10) or 50 Hz (n = 8) (Figure 2a) in line with our previous work [1, 10, 15]. Stimulation was delivered through each electrode consecutively, while recordings were simultaneously acquired from the remaining non-stimulating electrodes. Both the return and reference electrodes were placed subcutaneously. Therefore, none of the recorded signals were attributed to the reference electrode. None of the animals were allowed to recover from anesthesia at the end of the experiments and were humanely euthanized.

**Figure 1.**
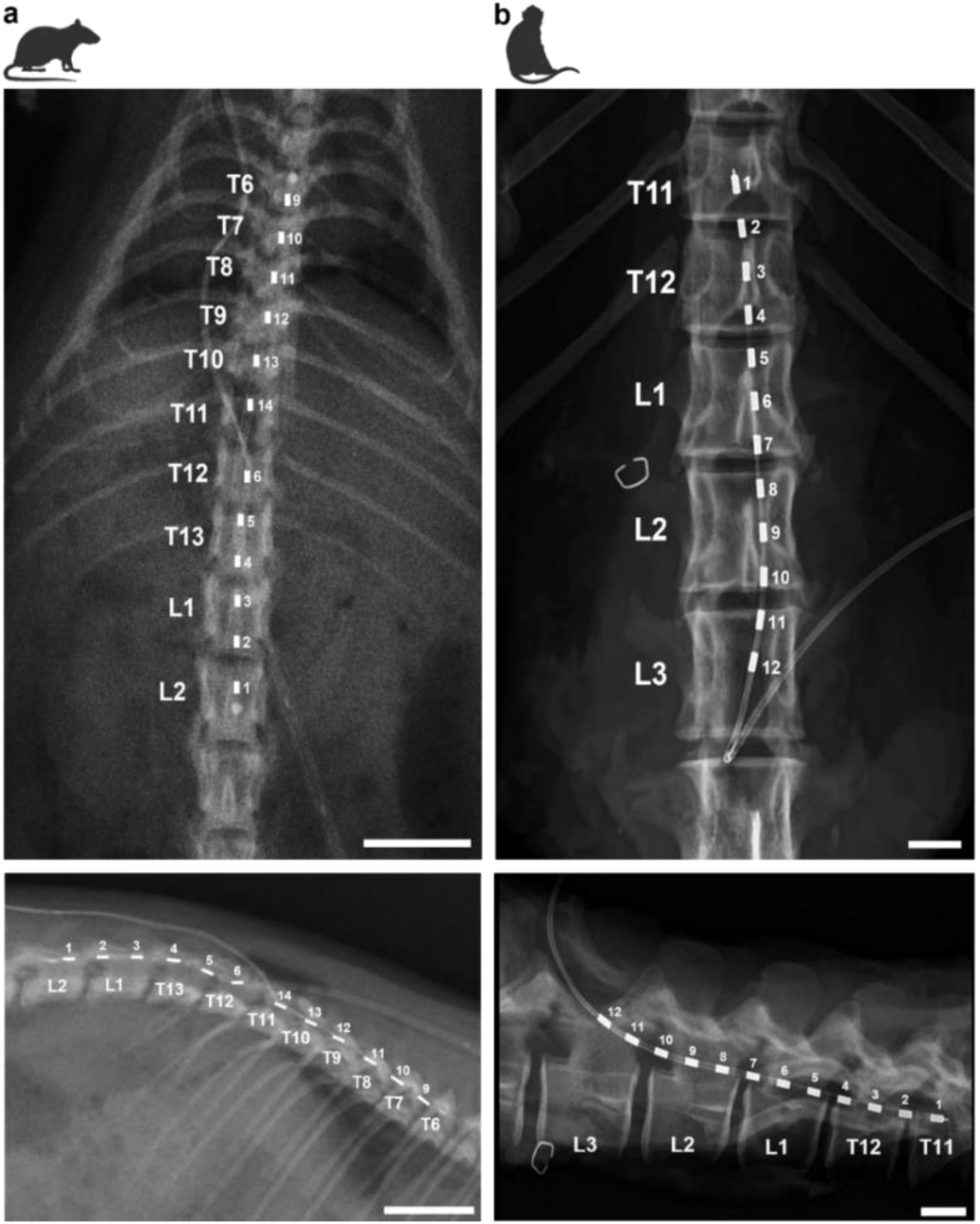
Representative lead positioning in the spinal cords of a rat and macaque. **a.** A representative dorsal X-ray image from a rat showing the placement of two epidural leads and confirming coverage from vertebral levels L2-T12 (lead 1) and T11-T6 (lead 2). Each electrode on the lead, along with the corresponding vertebral segments, is labelled (lead 1: 1-6, lead 2: 9-14). The representative lateral X-ray image is presented below. Note that the lumbosacral enlargement (spinal cord levels L1-S1) spans vertebral levels T12-L2, and that the spinal cord terminates at the intervertebral disc between the third and fourth lumbar vertebrae (∼L3-L4 vertebral level) [23, 24]. **b.** A representative dorsal X-ray image from a macaque showing the epidural lead covering the L3 to T11 vertebral levels. Each electrode and corresponding vertebral level are labelled (1-12). The representative lateral X-ray image is presented below. Note that the lumbosacral enlargement (spinal cord levels L1-S1) spans vertebral levels T12-L3, and that the conus medullaris terminates between L4 and S1 [23, 25, 26]. Scale bar: 10 mm.

**Figure 2.**
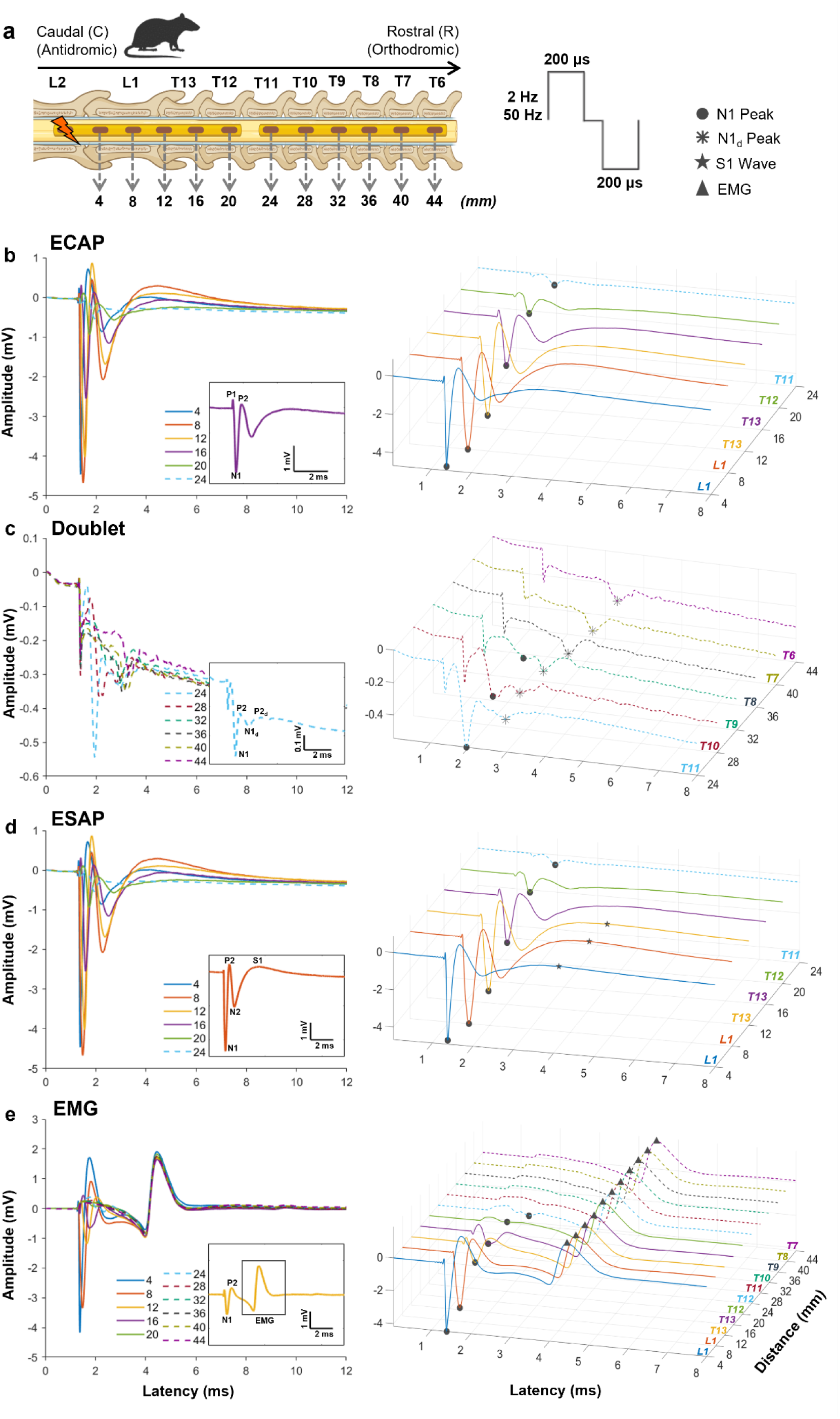
Epidural spinal electrophysiological signals evoked by spinal cord stimulation in rats. **a.** Schematic illustration of the experimental electrode setup, showing the distances (in mm) between the stimulating and recording electrodes, along with their corresponding vertebral level placements. Two leads were implanted epidurally: one caudally (T12-L2) and one rostrally (T11-T6). Stimulation was delivered through each electrode consecutively, while recordings were simultaneously acquired from the remaining electrodes. Return and reference electrodes were placed subcutaneously. Data presented in the figure refer to stimulation delivered through the most caudal electrode and recordings obtained from all remaining electrodes on both leads. Monopolar, biphasic stimulation was delivered in a stepwise manner (1 µA) with increasing current up to visible motor threshold, using a pulse width of 200 μs, at frequencies of 2 or 50 Hz. *Created with BioRender.com.* Evoked compound action potential (ECAP; **b**), doublet (**c**), evoked synaptic activity potential (ESAP; **d**), and electromyography (EMG) signals (**e**) recorded at distances of 4-24 mm (ECAP, ESAP, and EMG) or 24-44 mm (doublet) from the stimulating electrode. The N1 peaks are marked with dots (**b-e**), the secondary N1 peaks (N1d) with asterisks (**c**), the S1 waves with stars (**d**), and the EMG maxima with upward triangles (**e**). Each trace represents the average of two recordings at 0.23 mA (**b-d**) and fifty recordings at 0.30 mA (**e**), each evoked by separate stimulation pulses delivered at motor threshold within a 1-s period in a single rat. Insets highlight the characteristic morphology of each signal type. The solid line represents the lead implanted caudally, while the dotted line represents the lead implanted rostrally.

Recordings were averaged across approximately two traces at 2 Hz or fifty traces at 50 Hz per current level. Prior to analysis, the data were upsampled fivefold to improve temporal resolution. No offline filtering or artefact-specific correction procedures were applied. The amplifier was blanked during the stimulation to reduce the stimulus artifact [21]. Signal processing and characterization were performed using a custom-built analysis toolbox (MATLAB 2013 release; MathWorks, Inc.) along with custom scripts developed in MATLAB 2024.

#### Macaque

Anesthesia was initially induced with ketamine (10 mg/kg, i.m.), supplemented by medetomidine (3 μg/kg, i.m.) and midazolam (0.3 mg/kg, i.m.) or propofol (1.3-4.4 mg/kg, i.m.). General anesthesia was maintained using sevoflurane (2-3% in 100% oxygen) and alfentanil (0.4-0.57 μg/kg/min, i.v.). To enhance analgesia, meloxicam (0.3 mg/kg, i.m.) and paracetamol (25 mg/kg, i.m.) were administered. Antibiotic prophylaxis was provided with either cefotaxime (20 mg/kg, i.v.) or co-amoxiclav (12.5-20 mg/kg, i.v.), and methylprednisolone (5.4 mg/kg/h, i.v.) was given to minimize cerebral edema. Fluid balance was maintained with Hartmann’s solution (approximately 5 mL/kg/h, i.v.), with adjustments to maintain a total infusion rate of ∼10 mL/kg/h, accounting for concurrent drug administration. A tracheotomy was performed to enable positive pressure artificial ventilation. A central arterial line was inserted via the carotid artery for continuous blood pressure monitoring, and the bladder was catheterized. Body temperature was maintained using a thermostatically controlled heat pad and a warm air blanket. A laminectomy was performed to expose the cervical and lumbar spinal cord. Following this, sevoflurane was discontinued (reduced to 0%), and anesthesia was transitioned to continuous infusions of alfentanil (0.4-1.2 µg/kg/min, i.v.), ketamine (6-10 mg/kg/h, i.v.), and midazolam (0.3 mg/kg/h, i.v.), a regimen previously shown to support stable anesthesia while preserving central nervous system activity. Throughout the procedure, physiological parameters were closely monitored, including pulse oximetry, heart rate, arterial blood pressure, core and peripheral temperatures, and end-tidal CO₂. Rapid increases in heart rate or blood pressure in response to noxious stimuli, or gradual upward trends in these measures, were interpreted as signs of inadequate anesthesia, prompting supplemental doses of injectable agents as needed. Macaques were implanted with a twelve-electrode SCS lead (pellethane body with platinum-iridium electrodes; overall dimensions: 1.3 mm diameter; individual electrode size: 3.0 mm electrode length; designed and fabricated by Saluda Medical, Sydney, Australia) positioned in the dorsal epidural space. Electrodes were spaced 7 mm apart (center-to-center), spanning spinal vertebral levels T11 to L3 (Figure 3a). Lead placement was verified by X-ray imaging (Orange 1040HF, EcoRay, Seoul, South Korea) (Figure 1b). SCS was delivered using a clinical-grade neuromodulation system (Evoke System, Saluda Medical, Sydney, Australia). In line with clinically used settings and previous studies [16], and to reduce stimulation artefact as well as avoid the second cathode effect [22], tripolar, triphasic stimulation (with the active electrode located between two return electrodes) was delivered at 10 Hz with a 100 μs pulse width (Figure 3a). Tripolar, biphasic stimulation at 3 Hz with an 80 μs pulse width (Figure 3a) was used to detect ESAPs, following the methodology of Sharma et al. (2023), who reported that these signals exhibit greater stability at lower frequencies (1 Hz compared to 50 Hz) [15]. Stimulation was applied in a stepwise manner with increasing current up to 2-3 times the ECAP threshold and was delivered through each electrode consecutively, while recordings were simultaneously acquired from the remaining non-stimulating electrodes. The position of the return electrodes for each recording are indicated in Figure 3. Neuromuscular blockade (atracurium; 0.75 mg/kg/h, i.v.) was administered during the acquisition of all electrophysiological signals except EMG, to minimize movement artifacts. None of the animals were allowed to recover from anesthesia at the end of the experiments and were humanely euthanized.

**Figure 3.**
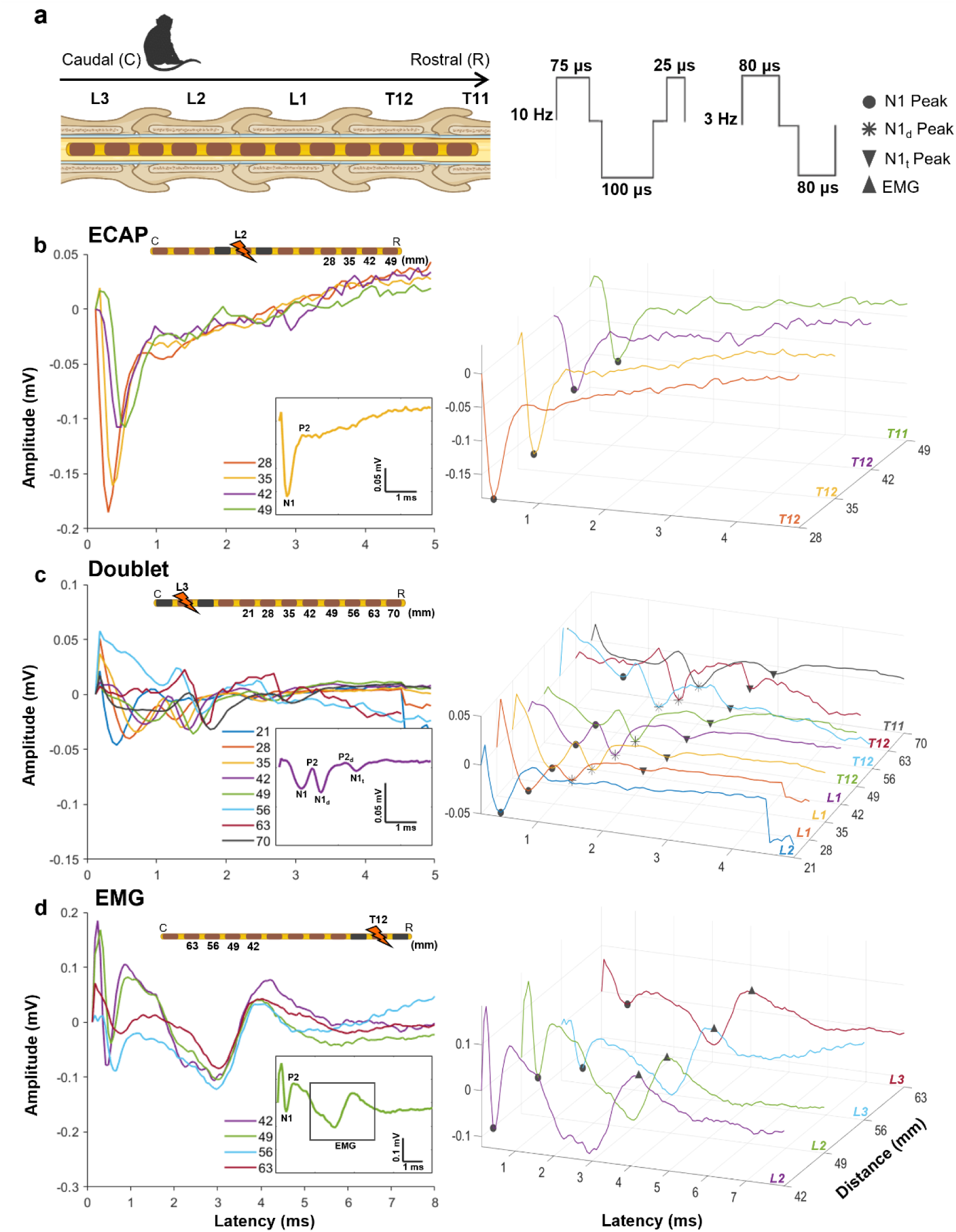
Epidural spinal electrophysiological signals evoked by spinal cord stimulation in macaques. **a.** Schematic illustration of the experimental electrode setup, showing the vertebral level placements. The lead was implanted epidurally (L3-T11). For the signals shown, stimulation at 1.5×Evoked Compound Action Potential (ECAP) threshold was delivered through different electrodes in a consecutive manner, while recordings were obtained from the remaining electrodes. The stimulation electrode is indicated for each signal in the schematic illustrations of the leads. Return electrodes are represented by black rectangles (as shown in **b-d**). Reference electrode was positioned on a second rostrally implanted lead at the furthest away contact. Stimulation was applied as tripolar, triphasic pulses at 10 Hz (100 μs) (**b, c**) or tripolar, biphasic pulses at 3 Hz (80 μs) (**d**). *Created with BioRender.com*. Evoked compound action potential (ECAP; **b**), doublet (**c**), and electromyography (EMG) signals (**d**) were recorded at distances of 28-49 mm (ECAP), 28-70 mm (doublet), and 42-63 mm (EMG) from the stimulating electrode. The N1 peaks are indicated by dots (**b-d**), the secondary N1 peaks (N1d) by asterisks (**c**), the tertiary N1 peaks by downward triangles (N1t) (**c**), and the EMG maxima by upward triangles (**d**). Each trace represents the average from a single macaque, recorded at a current of 0.75 mA (1.5×ECAPT) for panels **b-c**, and 1.7 mA for panel **d**. Insets emphasize the characteristic morphology of each signal type.

Recordings were averaged across approximately three traces at 3 Hz or ten traces at 10 Hz for each current level. Data were analyzed without upsampling. No offline filtering or artefact-specific correction procedures were applied. The amplifier was blanked during the stimulation to reduce the stimulus artifact [21]. Analyses were performed using a custom application (Plot Builder) in combination with custom scripts developed in MATLAB 2024 (MathWorks, Inc.).

In both species, the threshold for each electrophysiological signal was defined as the lowest stimulation current at which the signal could be reliably detected [1,10]. Signal amplitudes (mV) were calculated as the voltage difference between characteristic peaks: P2-N1 for ECAPs, P2_d_-N1_d_ for doublets (d refers to secondary ECAP), the maximum positive peak minus the preceding negative peak for EMG signals. For ESAPs, amplitude was defined as the voltage measured at the maximum point of the S1 peak. Conduction velocity (m/s) was estimated by measuring the latency of characteristic peaks, N1 for ECAPs and N1d for doublets, across spatially separated recording electrodes.

## RESULTS

### ECAP

ECAPs were reliably evoked and recorded across multiple segmental levels of the spinal cord during SCS in both rats (L1-T8) and macaques (L2-T11). As the earliest electrophysiological responses detected in epidural spinal recordings, ECAPs were distinguished by their short latencies relative to other spinal signals described in this study (Figures 2b and 3b). Morphologically, ECAPs exhibited a characteristic triphasic waveform, consisting of an initial positive peak (P1), followed by a negative peak (N1), and a second positive peak (P2), with a total duration of approximately 1 ms from P1 to P2 (Figures 2b and 3b). The ECAP threshold was 32.00 ± 3.10 µA in rats (n = 18 out of 18) and 375.00 ± 25.00 µA in macaques (n = 2 out of 2).

As illustrated in Figure 4a for rats, ECAP amplitude increased linearly with stimulation intensity following suprathreshold activation, up to the motor threshold. Although the waveform remained consistent across recording sites (L1-T8), electrodes located farther from the stimulation source exhibited longer latencies and reduced amplitudes (Figure 2b and 2c). Comparable results were observed in macaques (Figure 3b, 3c and 5a). Average conduction velocities were 30.70 ± 1.81 m/s in rats (n = 18) (Figure 4b) and 82.75 ± 2.46 m/s in macaques (n = 2) (Figure 5b).

**Figure 4.**
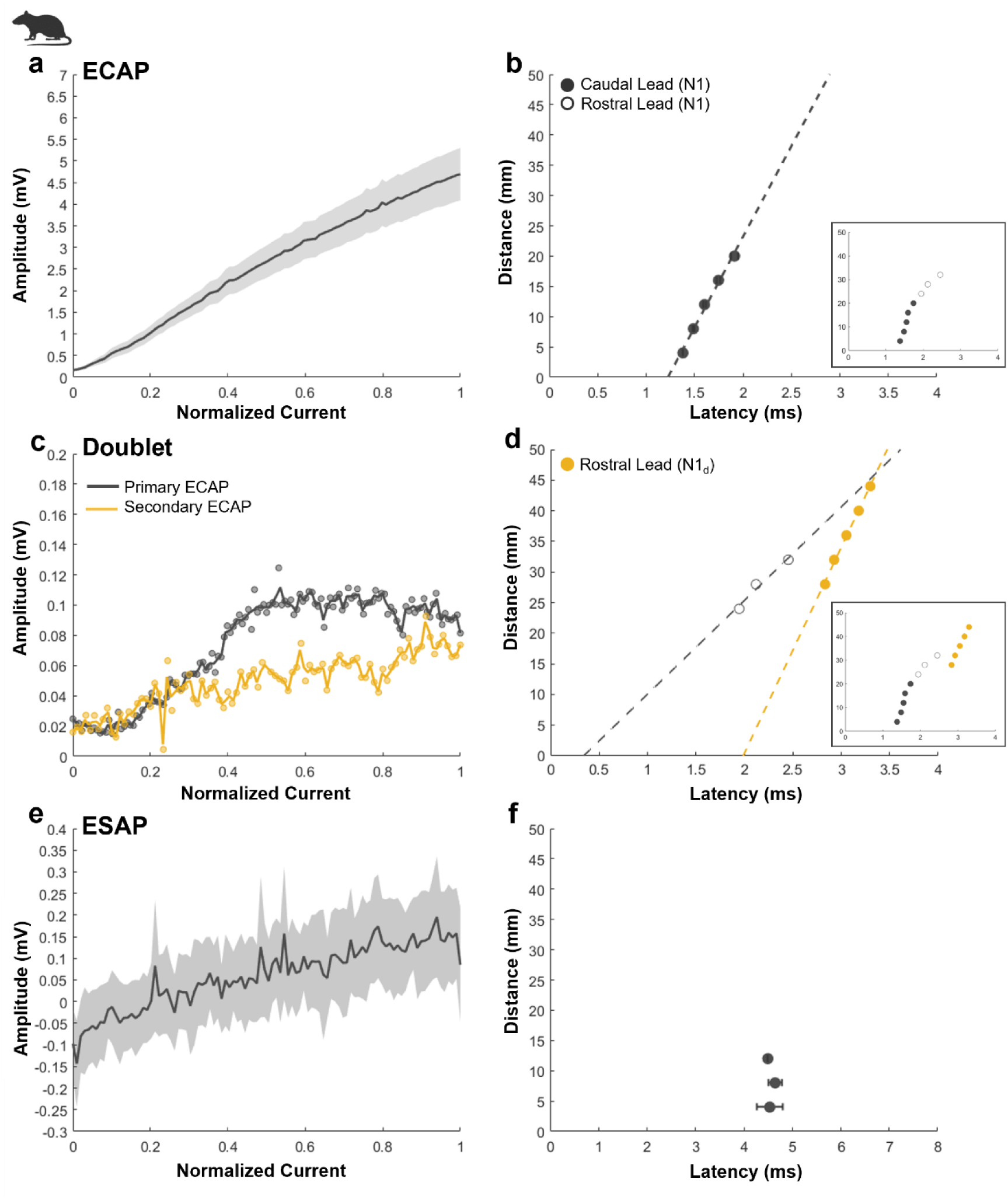
Spinal cord stimulation (SCS)-evoked neural responses in rats. **a.** Mean evoked compound action potential (ECAP) amplitude as a function of stimulation current during 2 Hz and 50 Hz SCS recorded orthodromically 4 mm from the stimulation site (n = 18). **b.** Mean ECAP distance vs. latency during 2 Hz and 50 Hz SCS across rats (n = 18; mean. 30.70 ± 1.81 m/s) calculated from N1 peaks. The inset shows ECAP distance vs. latency from a rat exhibiting doublets in panel c. Filled circles represent N1 latencies calculated from the caudally implanted lead, whereas open circles represent N1 latencies calculated from the rostrally implanted lead. The slope of the linear fit indicates the conduction velocity. **c.** Doublet amplitude vs. stimulation current during 2 Hz SCS (n = 1), recorded orthodromically ∼28 mm from stimulation site. Gray shows primary ECAP, and yellow shows secondary ECAP. **d.** Doublet distance vs. latency during 2 Hz SCS (n = 1). The slope of the linear fits indicates the conduction velocities. Primary and secondary ECAP conduction velocities were 15.28 and 33.62 m/s, respectively. Primary ECAP conduction velocity was calculated from N1 peaks, while secondary ECAP conduction velocity was calculated from N1d peaks. In the inset, filled circles represent N1 latencies calculated from the caudally implanted lead, open circles represent N1 latencies calculated from the rostrally implanted lead, whereas yellow filled circles represent N1d latencies recorded from the rostrally implanted lead. **e**. Mean evoked synaptic activity potential (ESAP) amplitude vs. stimulation current during 2 Hz and 50 Hz SCS across rats (n = 4), recorded orthodromically 8 mm from stimulation site. **f.** Mean ESAP distance vs. latency during 2 Hz and 50 Hz SCS across rats (n = 4), recorded orthodromically. All data were collected with 200 μs pulse width. Stimulation current was normalized between the relevant signal threshold (0) and the motor threshold (1) (**a, c, e**). Distance vs. latency were plotted and conduction velocity was calculated at motor threshold. Standard error of the mean (± SEM) is indicated by the shaded region (**a**, **e**) and error bars (**b**, **f**).

**Figure 5.**
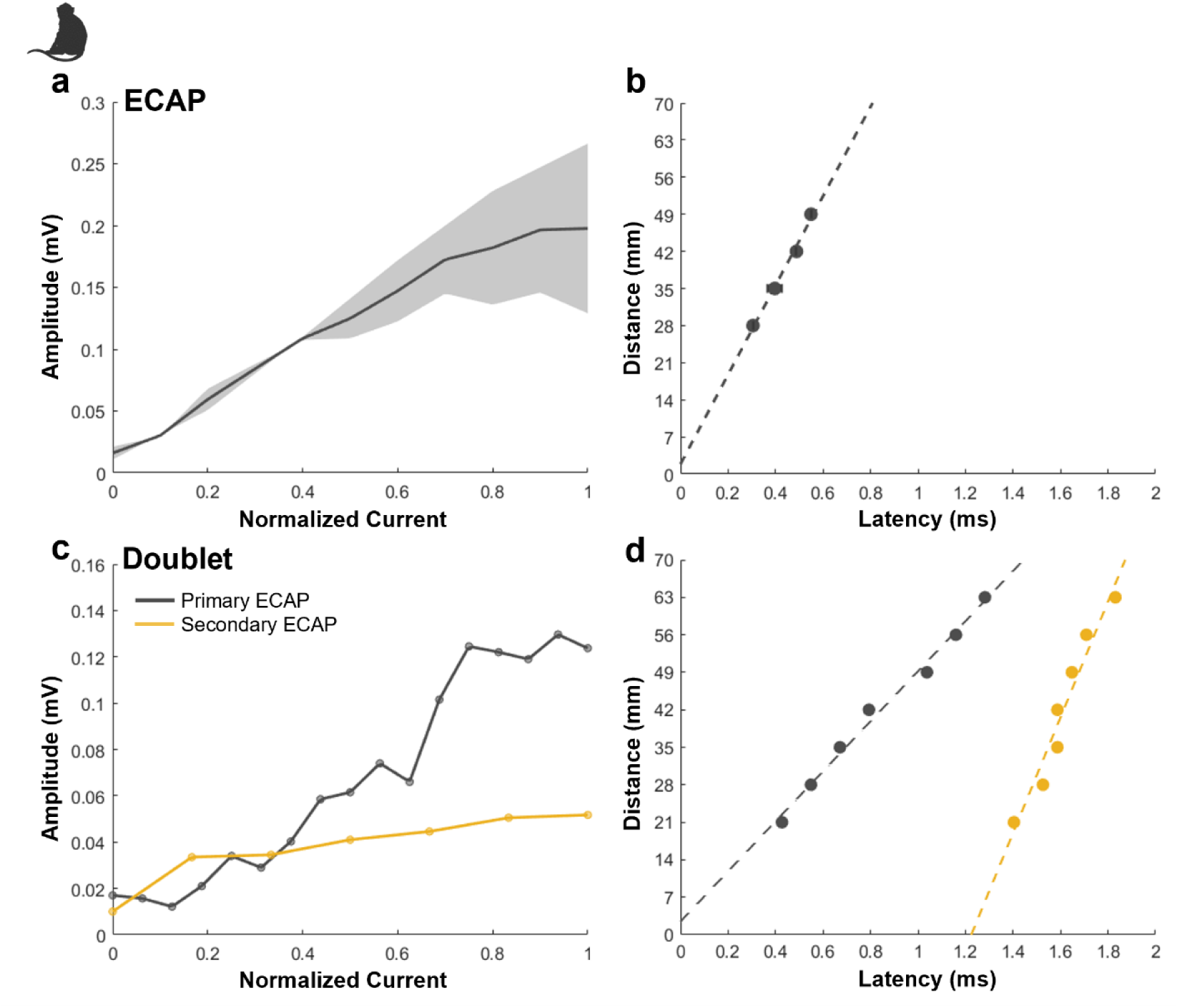
Spinal cord stimulation (SCS)-evoked neural responses in macaques. **a.** Mean evoked compound action potential (ECAP) amplitude as a function of stimulation current during 10 Hz SCS (n = 2), recorded orthodromically from the electrode 28 mm away from stimulation site. Stimulation current was normalized between ECAP threshold (0) and 2×ECAP threshold (1). **b.** Mean ECAP distance vs. latency during 10 Hz SCS across macaques (n = 2; mean. 82.75 ± 2.46 m/s), calculated from N1 peaks. Gray dots and dashed line show group mean. The slope of the linear fit indicates the conduction velocity. **c.** Doublet amplitude vs. stimulation current during 10 Hz SCS (n = 1), recorded orthodromically 28 mm from stimulation site. Gray shows primary ECAP data, and yellow shows secondary ECAP data. Stimulation current was normalized between the relevant signal threshold (0) and 3×ECAP threshold (1). **d.** Doublet distance vs. latency during 10 Hz SCS (n = 1). The slope of the linear fits indicates the conduction velocities. Primary and secondary ECAP conduction velocities were 46.74 and 108.07 m/s, respectively. Primary ECAP conduction velocity was calculated from N1 peaks, while secondary ECAP conduction velocity was calculated from N1d peaks. All data were collected with 100 μs (**a-d**) pulse width. Standard error of the mean (± SEM) is indicated by the shaded region (**a**).

### Doublet

Doublets appeared as complex signals characterized by multiple negative peaks, two in one of the rats (out of 18) and three in both macaques, representing two (or three) separate ECAPs. In Figure 2c distinct primary (N1) and secondary (N1d) peaks are observed at levels T11 to T8, whereas beyond T8 only the continuation/propagation of N1d peak was observable and the primary N1 peak could no longer be observed. In Figure 3c, N1 and N1d peaks are clearly distinguishable at levels L2 to T12, but only N1d peaks are observable at T12 to T11 (distances of 63 and 70 mm away from stimulation site). The secondary ECAP, elicited 1.65 ms in the rat and 1.36 ms in the macaque after the primary ECAP was reliably detected in both species (Figures 2c and 3c; asterisk). In macaques, a tertiary ECAP was also observed, emerging around 2 ms after the primary ECAP and 1 ms after the secondary ECAP (Figure 3c; downward triangle). In one macaque, secondary and tertiary ECAPs were observed at a distance between 28 and 70 mm away from stimulation site (Figure 3c). In the second macaque, secondary ECAPs were also observed between 28 and 70 mm away, however tertiary ECAPs were only observed beyond approximately 77 mm away from stimulation (data not shown).

Doublets were only evoked when stimulation was applied to the lower thoracic to upper lumbar spinal regions (shown in Figure 2c as L2 and Figure 3c as L3). They were recorded exclusively in the orthodromic direction (rostral), predominantly within the lower to upper thoracic spinal cord (T11-T6 in Figure 2c), though they were also detected in the upper lumbar region in the macaques (L1-T11 in Figure 3c). Both the secondary and tertiary ECAPs retained the triphasic morphology characteristic of the primary ECAP. The threshold for secondary ECAP was 143 µA (n = 1) in the rat, corresponding to a primary ECAP threshold of 105 µA (n = 1), recorded on the same electrode. In macaques, secondary ECAP thresholds were 550 µA (n = 1) and 950 µA (n = 1), while primary ECAP thresholds on the same electrode were 550 µA (n = 1) and 400 µA (n = 1), respectively. In the macaque exhibiting tertiary ECAPs at 35 mm, the threshold was approximately 700 µA (n = 1).

Their amplitudes (measured as P2_d_-N1_d_ in mV for the secondary, and P2_d_-N1t in mV for the tertiary; t refers to tertiary ECAP) increased in response to rising stimulation current (Figure 4c and 5c). Unlike the primary ECAP, the amplitudes of these later components increased with recording distance (Figures 2c and 3c). Similar to the primary ECAP, both secondary and tertiary components demonstrated clear propagating behavior, with latencies increasing at electrodes positioned farther from the stimulation site (Figure 4d and 5d). Conduction velocity of the secondary ECAP was 33.62 m/s (n = 1) in the rat, while primary ECAP conduction velocity was 15.28 m/s (n = 1) (Figure 4d). In macaques, secondary ECAP conduction velocities measured 81.54 m/s and 108.07 m/s, with corresponding primary ECAP velocities of 54.81 m/s and 46.74 m/s (n = 2) (Figure 5d). In one of the macaques, tertiary ECAPs exhibited a conduction velocity of 79.49 m/s (n = 1) (calculated from N1t peaks).

### ESAP

ESAPs were detectable in rats at 2 and 50 Hz, (Figure 2d; star; n = 4 out of 18) but were not observed in macaques. Morphologically, ESAPs were defined by a positive S1 wave that appeared at a longer latency (∼3-4 ms) following a secondary negative peak (N2) in the epidural recordings. The S1 wave exhibited a prolonged duration of approximately 6 ms, in contrast to the ∼1 ms duration of ECAPs. The average ESAP threshold was 91.00 ± 12.00 µA (n = 4).

Similar to ECAPs, the S1 wave amplitude increased in response to rising stimulation current (Figure 4e). However, unlike ECAPs, ESAP amplitudes did not consistently decrease with increasing distance from the stimulation site. Instead, ESAPs exhibited a spatially localized maximum, with the largest signals consistently recorded at a specific electrode (e.g., electrode 3 located 8 mm from the stimulation site at L2; Figure 2d). ESAPs were only detectable when the recording contact was positioned over the L1/T13 vertebrae, with peak responses consistently observed at the L1 level, suggesting that anatomical specificity may be a defining characteristic of ESAPs. Furthermore, ESAP latencies did not increase with distance from the stimulation site, consistent with their non-propagating nature (Figure 4f).

### EMG

EMG signals were recorded in one representative animal for each species. They were detected in epidural recordings and were distinct from ECAPs, doublets and ESAPs, in morphology and stimulus-response characteristics. EMG signals appeared as either biphasic or triphasic waveforms 3-5 ms after the stimulus, which is approximately 2-4 ms after the primary ECAP (Figure 2e and 3d). These signals had a slightly longer duration than ECAPs (∼2 ms).

Similar to ECAPs, EMGs were evoked and recorded across multiple spinal levels in the rat (L1-T7; Figure 2e) and the macaque (L2-L3; Figure 3d). The EMG threshold was 245 µA in the rat (n = 1) and 900 µA in the macaque (n = 1).

A key distinguishing feature of EMG signals was their step-like recruitment pattern, characterized by the abrupt appearance of EMG responses once the stimulation threshold was reached, rather than a gradual increase with stimulation intensity. In both species, EMG amplitudes were consistent across recording sites, showing no change in amplitude (in channels where the signal was detected) as the distance from the stimulation site increased (Figure 2e and 3d). Unlike ECAPs and doublets, EMGs were non-propagating, meaning their latency remained constant regardless of the distance of the recording electrode from the stimulation site.

## DISCUSSION

We present here a summary of the distinct electrophysiological signals evoked and recorded epidurally during SCS, each distinguishable by unique waveform features and stimulus-response characteristics, including latency and amplitude. This study highlights their defining properties and provides representative examples to facilitate interpretation (Table 1). By incorporating data from both rats and macaques, we enable cross-species comparisons that may offer translational insights. Below, we further discuss the relevance of each signal to spinal processing. Unlike ECAPs, other evoked signals may be affected by aberrant spinal circuitry, particularly in maladaptive states such as neuropathic pain, and thus may offer a deeper understanding not only of the mechanisms underlying SCS but also of the pathophysiology of chronic pain.

**Table 1.**
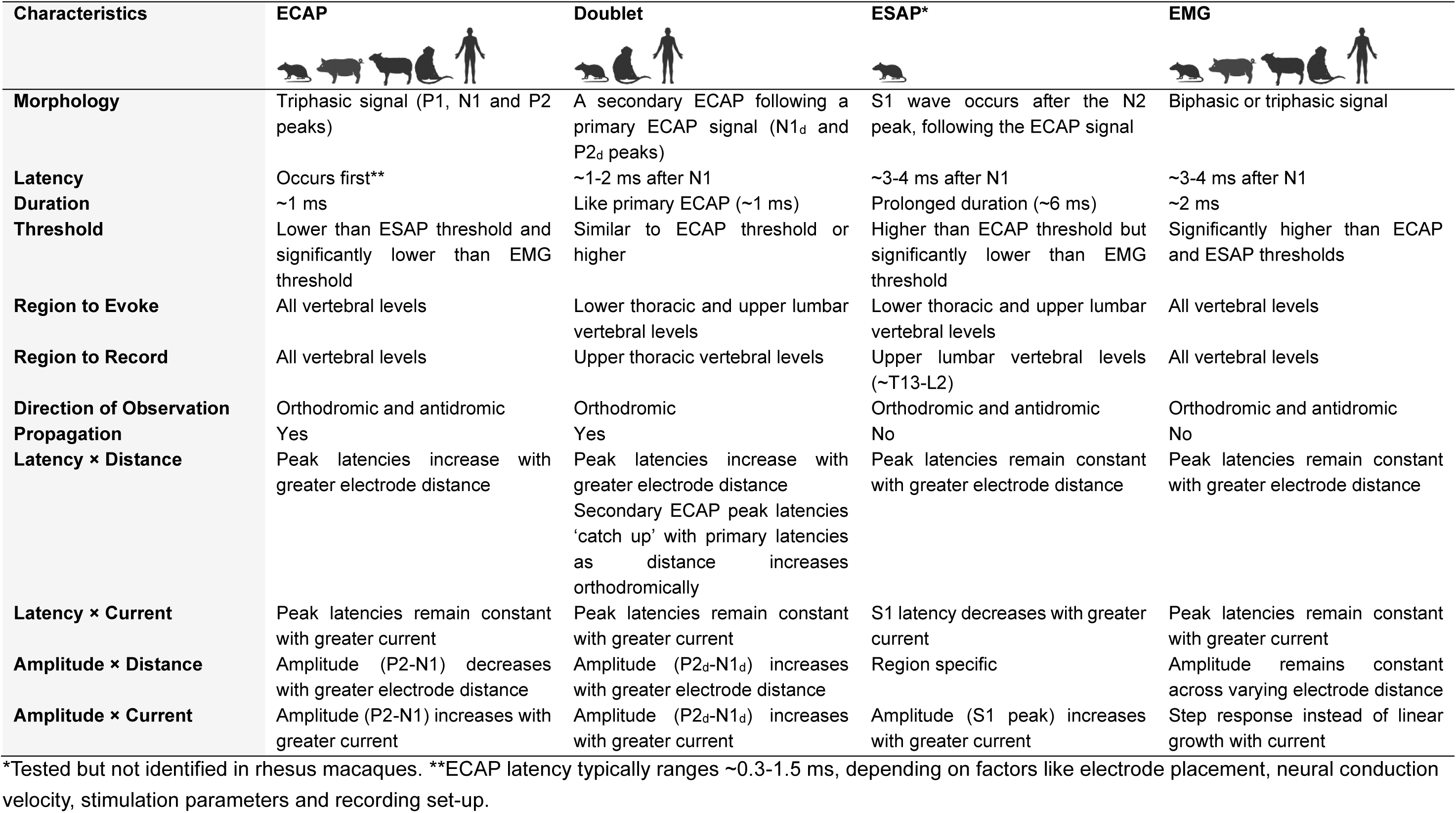
A summary of the defining characteristics of electrophysiological signals detected in epidural spinal recordings, including evoked compound action potentials (ECAPs), doublets, evoked synaptic activity potentials (ESAPs), and electromyographic (EMG) signals, as identified in current and previous studies, with species indicated by the corresponding icons.

### ECAP

ECAPs recorded in this study are consistent with those reported previously across multiple species, including rodents, large animals (e.g., sheep and pig), non-human primates and humans. The characteristic triphasic morphology (P1-N1-P2) observed here aligns with typical extracellular recordings of axonal CAPs, where the P1 peak reflects the initial depolarization, the N1 valley corresponds to axonal depolarization driven by Na⁺ influx, and the P2 peak reflects the subsequent hyperpolarization driven by K⁺ efflux [2]. This morphology is conserved across species, supporting the cross-species reliability of ECAP recordings [2].

Consistent with previous studies, ECAPs recorded during SCS primarily reflect the activation of fast-conducting DC fibers, specifically large-diameter myelinated Aβ afferents, as confirmed by conduction velocity measurements [1–5, 14]. A novel observation in this study is the apparent slowing of ECAP conduction velocity with increasing distances, observed in both rats (Figure 4b and 4d) and macaques (Figure 5b and 5d). Similar findings have been reported in anaesthetized cats, where the conduction velocity of group I and II afferent fibers progressively decreased as they ascended the DC [27]. This deceleration may indicate a greater relative contribution of smaller-diameter fibers to the ECAP as it propagates rostrally. One possible explanation for this deceleration is that larger-diameter axons terminate at lower spinal levels; less than 25% of lumbar DC axons reach the brainstem [28]. Supporting this, a study by Yamamoto et al. (1996) in rats demonstrated that only 50.1% of axons present in the DC at the L3 level ascend to the T6 level, with most large-diameter fibers terminating or thinning out between these regions [29]. Further investigation is needed to confirm these findings and further characterize this phenomenon in human subjects.

In addition to conduction velocity, changes in ECAP morphology, such as amplitude and width, may provide insight into DC fiber dynamics and their modulation by SCS parameters. For example, high-frequency SCS at 200, 500, and 1000 Hz reduced ECAP amplitude and increased peak latencies and ECAP width, effects likely reflecting asynchronous firing of DC axons [14]. These findings highlight ECAPs as an indicator of DC activation and support their utility in advancing both the mechanistic understanding and therapeutic optimization of SCS.

### Doublet

Here, we observed secondary ECAPs in both rodents and macaques that share core characteristics with previously described PSDC signals in humans [16]. Additionally, in macaques, we identified a tertiary ECAP which exhibited characteristics closely resembling those of secondary ECAPs, albeit occurring at a later latency. Maruyama et al. (1982) have previously described a similar signal recorded with epidural electrodes in response to cauda equina stimulation in humans [30].

Doublets reported in this study were exclusively evoked when stimulating at lower vertebral levels and recorded at upper levels, in the orthodromic (rostral) direction. These responses exhibited higher conduction velocities compared to the N1 component of primary ECAPs, consistent with previous findings in humans [16]. The presence of these additional negative peaks suggests activation of spinal projection neurons, which are likely recruited following electrical stimulation of DC axons and may represent output from endogenous spinal processing. The administration of a neuromuscular blocker in the macaques used when obtaining all recordings except from the EMG signal in this study, effectively rules out any potential contamination of the signals by myogenic activity. Moreover, our interpretation aligns with previous studies reporting PSDC neurons to be predominantly found in the lumbar and cervical enlargements in animals [31, 32]. While doublets described here are consistent with the hypothesized activation of the PSDC pathway, contributions from alternative spinal pathways involving postsynaptic activation, such as the spinothalamic tract [17], cannot be ruled out, particularly given that doublets exhibit higher activation thresholds compared to primary ECAPs.

Gmel et al. (2023) found no evidence that secondary ECAP signals in humans resulted from direct activation of nerve rootlets, based on propagation times for the secondary ECAP [16]. A comparable calculation, extrapolating the two trendlines to their y-intercepts, shows that in the rat the intercepts differ by 61.6 mm. This would require stimulation pulses to activate a fiber bundle located 6.16 cm away, without affecting any intervening fibers. In the monkey, the intercepts are 134 mm apart, implying a required activation distance of 13.4 cm under the same unlikely condition. Furthermore, care must be taken with stimulating contact configurations (cathodes and anodes) and charge delivery, since second cathode effects can manifest as multimodal signals [19, 22].

### ESAP

First identified in epidural lead recordings by Sharma et al. (2023) in rat, the ESAPs are reproduced here also in rats, consistent with those previously described [15]. Features of the S1 component of the ESAP are reported to be consistent with intracellularly recorded monosynaptic responses of dorsal horn neurons in *in vitro* models following DC stimulation [19]. Although not demonstrated in the present study, the pharmacological sensitivity of ESAPs to AMPA receptor blockade provides strong evidence that they originate from synaptic, rather than axonal, activity [15]. Sharma et al. further verified that the S1-wave was not a stimulation artifact and/or a reflection of hindlimb/trunk EMG [15]. Importantly, the spatial specificity of ESAP detection, unlike the widespread propagating nature of ECAPs, suggest that ESAPs reflect local connectivity (circuits) (Table 1) [15]. Additionally, Sharma et al. demonstrated that ESAPs, but not ECAPs, exhibited gradual adaptation in response to 50 Hz stimulation [15]. This habituation is consistent with, though not exclusive to, synaptic processing mechanisms and mirrors findings from superficial dorsal horn interneuron responses to 50 Hz SCS [15, 33–35]. Together, these observations support the hypothesis that ESAPs may serve as an epidurally measurable marker of dorsal horn activity in rodents.

### EMG

Building on the identification of neural signals such as ECAPs, doublets and ESAPS, it is also essential to distinguish EMG signals which can be detected via the epidural lead. EMGs are often detected within the same latency as the other neural responses and can introduce a source of signal contamination, especially when interpreting overlapping components [19]. However, the non-propagating nature of EMGs, along with their step-like appearance, higher activation thresholds and consistent amplitude across recording sites, help differentiate them from neural signals such as ECAPs, doublets and ESAPs.

EMG signals can be observed across multiple species, including rodents, large animals (e.g., sheep and pig), non-human primates and humans, confirming their relevance in both preclinical and clinical contexts. In our study, EMG signals were recorded in only one representative animal per species, as our original aim was not to focus on these signals. In anesthetized rats, a visible motor response, used to determine the point at which current increase was stopped, typically occurs before EMG activity becomes detectable in epidural recordings; therefore, EMG was not recorded in additional rats. In the single macaque, EMG signals were obtained once the effects of the neuromuscular blocker had begun to wear off.

It remains to be confirmed whether EMG signals induced by SCS result from direct activation of motor neuron axons, as they exit the anterior spinal cord via the ventral roots, or from monosynaptic activation of motor neurons following stimulation of proprioceptive afferents in the dorsal roots. In peripheral nerve studies, the H-wave typically has a threshold lower than the M-wave, reflecting the greater excitability of afferent fibers compared to efferent fibers [36]. Given the dorsal roots’ closer proximity to the stimulating electrodes relative to motor axons in the ventral roots, SCS-induced EMG signals in humans and large animal models are most likely reflex-mediated. In rodents, however, the deep DCs contain descending corticospinal fibers that may be directly activated at higher SCS intensities, subsequently activating motoneurons [37]. Thus, similar to ESAPs and potentially doublets, EMG signals likely represent an output of spinal processing.

### Limitations

There are several limitations in the current study that can be addressed by future research. First, the electrophysiological signals reported here were recorded in preclinical SCS models without experimentally induced neuropathic pain. As described above, ESAPs, EMG signals, and potentially doublets, reflect outputs of spinal processing detectable via epidural recordings, and may therefore provide insights into the pathophysiology of conditions involving disrupted spinal function. For example, diabetic neuropathic pain may arise from spinal cord disinhibition [38]. Future studies should aim to characterize the presence and properties of these electrophysiological signals under neuropathic pain conditions.

Second, ESAPs were detected in a small proportion of rats and a small number of doublets were detected in one rat and both macaques. Although our methodology followed that of Sharma et al. (2023), their experimental design and surgical approach was different (e.g., multi-level laminectomy, different anesthetic regimen, and female rats), which may explain their ability to detect ESAPs in a greater proportion of rats. The absence of ESAPs in our macaque model may reflect species-dependent differences in spinal processing, limitations in recording sensitivity (e.g., CSF thickness separating the dorsal horn from recording electrodes), or differences in the stimulation frequency and the multimodal anesthetic regimen, which could affect synaptic transmission. Pilot studies in other large animals (sheep and bovine) suggest slow waves of <50 µV (or 0.1×ECAP), highlighting the need for an enhanced signal-to-noise ratio. Similarly, doublets may be influenced by the same factors, highlighting the need for further investigation into the conditions required to detect ESAPs and doublets more consistently, including higher-order species. Notably, slow-wave signals (typically ∼50 µV when recorded epidurally) were consistently observed in human spinal cord neurophysiology studies [30, 33, 34] and indeed formed the foundation for the development of gate-control theory and, ultimately, SCS [35, 39]. As such, slow waves are a common signal in human spinal electrophysiology, although they have yet to be identified in human SCS epidural lead recordings.

Third, all recordings in this study were performed under anesthesia to ensure a controlled and stable experimental environment. Our unpublished observations indicate that anesthesia affects ECAP thresholds and latencies. However, whether the characteristics of the other signals remain consistent with those observed under anesthesia is still unknown. Future studies in awake animals will be needed to address this question. Lastly, we acknowledge that all animals in our study were male. Future research should include female subjects to determine whether sex-based differences influence the characteristics of these spinal signals.

## CONCLUSIONS

Epidural spinal recordings, including ECAPs, doublets, ESAPs, and EMGs provide insights into the neural processes engaged by SCS. This study summarizes key characteristics of these signals, as outlined in Table 1, to support their differentiation and classification in future studies based on waveforms, recruitment thresholds, and physiological origins. Continued investigation into the origins, functional relevance, and clinical significance of these signals may provide critical insights into SCS mechanisms, inform therapy optimization, and guide the development of more effective SCS interventions. Species-specific patterns, potentially reflecting neuroanatomical or technical differences that influence signal detectability in preparations of varying sizes, emphasize the importance of cross-species analyses to improve the translational relevance of neuromodulation research.

### Data statement

The datasets used and/or analyzed during the current study are available from the corresponding author on reasonable request.

### Declaration of generative AI and AI-assisted technologies in the writing process

During the preparation of this work, the authors used ChatGPT in order to improve the readability and language of the manuscript. After using this tool, the authors reviewed and edited the content as needed and take full responsibility for the content of the publication.

## Glossary

ARRIVE: Animal Research: Reporting in *In Vivo* Experiments
AWERB: Animal Welfare Ethical Review Body
CAP: Compound Action Potential
DC: Dorsal Column
ECAP: Evoked Compound Action Potential
ECAPT: Evoked Compound Action Potential Threshold
EMG: Electromyography
ESAP: Evoked Synaptic Activity Potential
N1: First Negative Peak
P2: Second Positive Peak
SCS: Spinal Cord Stimulation
SEM: Standard Error of the Mean

